# Early life adversity and White Matter Microstructural Organization - a systematic review

**DOI:** 10.1101/2025.02.04.634280

**Authors:** Orla Mitchell, Darren W Roddy, Michael Connaughton

**Affiliations:** Department of Psychiatry, Royal College of Surgeons in Ireland, Dublin 2, Ireland

## Abstract

Early life adversity, defined as exposure to stressful events during childhood, is a significant risk factor for the development of psychiatric disorders. Diffusion tensor imaging studies employing tract-based spatial statistics have shown microstructural abnormalities in white matter among individuals exposed to early life adversity; however, robust conclusions are yet to be drawn. This systematic review synthesizes findings of previous tract-based spatial statistics studies to identify the white matter alterations in adult brains exposed to early life adversity, in papers with methodological consistency. The literature search (April 2024) was conducted to identify tract-based spatial statistics studies that compared diffusion metrics between adults exposed to early life adversity and adults not. Embase, Pubmed, and PsycInfo were searched, retrieving 2458 articles. Following deduplication, 1739 titles and/or abstracts were screened. 1699 articles were excluded, and 40 full texts were reviewed. Seven articles, reporting on 764 subjects, met the inclusion criteria and were included in the narrative synthesis. Compared to controls, adults exposed to early life adversity showed lower fractional anisotropy values in white matter tracts of the limbic and visual processing systems, specifically the anterior thalamic radiation, inferior longitudinal fasciculus, corona radiata, uncinate fasciculus, inferior fronto-occipital fasciculus, and cingulum bundle. This systematic review highlights that early life adversity may underlie emotional dysregulation and contribute to an increased risk of psychopathology in later life and explores the potential neurobiological mechanisms that underpin these structural changes. Understanding these associations is crucial for developing targeted interventions aimed at mitigating the long-term impact of early life adversity.

## 1. Background

Early life adversity (ELA) refers to the exposure to stressful or traumatic experiences during childhood or adolescence, which represent a deviation from the expected environment (Duffy et al., 2018). Given that the expected environment encompasses inputs necessary for normal brain development, instances of ELA often necessitate substantial psychological, behavioral, or neurobiological adaptations by the exposed child (McLaughlin, 2016). Exposure to these stressors can occur in the context of threat or deprivation. Threat refers to direct harm or risk of harm, like abuse or exposure to violence, and deprivation refers to the lack of care, including lack of adequate caretaking, cognitive or social stimulation and extends to unpredictability. Exposure to ELA is common, with a 2018 study revealing that 61.55% of the U.S. population reported at least one adverse childhood experience, and 24.64% three or more (Merrick et al., 2018). Previous research has found strong associations between ELA and negative physical and mental health outcomes in adulthood, as well as engagement in health harming and negative relational behaviors (Hughes et al., 2017). Particularly, in mental health research, ELA has been implicated in the development of major depressive disorder, schizophrenia, and personality disorder (Coughlan & Cannon, 2017; Lenze et al., 2008; Pietrek et al., 2013; Zhang et al., 2020).

The mechanism by which ELA may elevate the risk of adverse outcomes remains poorly understood, but existing literature suggests it may involve a series of neurobiological adaptations resulting in alterations in brain structure, function and connectivity (Ancelin et al., 2021; Callaghan & Tottenham, 2016; Hakamata et al., 2022). Given that in early life, the brain is developing and has periods of heightened white matter plasticity (McLaughlin, 2016; McLaughlin et al., 2014), research suggests that ELA may be particularly impactful on white matter development (McManus et al., 2022). These white matter microstructural changes identified in association with exposure to ELA may be due to mechanisms related demyelination, such as vulnerability to oxidative stress and proinflammatory processes (Sapolsky et al., 2000; Schiavone et al., 2012). White matter is the tissue that joins different brain regions and is essential for maintaining typical brain function and cognition (Fields, 2010). Optimal white matter microstructural organization is necessary for fast and efficient processing and optimal brain functioning (Penke et al., 2010). Disruptions to white matter organization may impact processing speed (Kennedy & Raz, 2009), memory (Wannan et al., 2022), and ultimately has been linked to psychopathology (Murphy & Frodl, 2011; Vitolo et al., 2017), with a recent paper identifying that white matter microstructure across the brain is related to general psychopathology and this is not only with regards to specific tracts (Neumann et al., 2020). Exposure to stress in early life has even been linked to lasting alterations in white matter microstructure that persist into adulthood (Lu et al., 2013; McCarthy-Jones et al., 2018), however, robust patterns or conclusions have not yet been drawn.

Diffusion Weighted Imaging (DWI), an MRI acquisition technique, provides an understanding of brain microstructure through differences in the Brownian motion of water molecules (Baliyan et al., 2016). Diffusion Tensor Imaging (DTI), a method of modelling of DWI data (Soares et al., 2013), provides information regarding the orientation and quantifies the strength of directionality of diffusion. Importantly, DTI metrics can offer clinically relevant insights into the microstructural organization of white matter. Fractional anisotropy (FA), a widely used DTI metric, summarizes white matter microstructural organization. A reduction in FA values can indicate processes such as demyelination, axonal damage, or a decline in white matter coherence (Van Hecke et al., 2015; Zhang & Burock, 2020). Tract-Based Spatial Statistics (TBSS) is a widely used DTI analysis technique for studying white matter microstructural organization (Smith et al., 2006). TBSS is an automated, observer-independent technique that conducts voxel-wise comparisons of white matter diffusion properties (Smith et al., 2006). This method offers several advantages, including reliable registration of subjects’ FA data to a common space by projecting it onto a mean FA tract skeleton and implementation of permutation statistics that do not rely on the normal distribution of data (Smith et al., 2006). Unlike other methods, it avoids the use of a smoothing kernel and minimizes partial volume effects (Smith et al., 2006).

TBSS analysis of DTI data is one of the most popular and commonly published in the diffusion imaging field (Bach et al., 2014; Raffelt et al., 2015). Synthesizing the white matter changes associated with ELA across methodologically consistent studies may provide vital insights into the underlying neurobiological mechanisms of ELA and its role in the development of psychopathology. This systematic review is novel in its focus on the long-term effects of ELA on white matter microstructure, rather than in children or adolescents. By exploring white matter microstructural changes identified in adult brains, this review offers insights into whether alterations in brain structure persist beyond periods of high developmental plasticity, potentially explaining the mental and physical health risks observed in those exposed to ELA. Additionally, by excluding studies on participants with psychiatric diagnoses, this review aims to isolate white matter changes specific to ELA, enhancing understanding of its direct impacts and potentially identifying structural biomarkers that may contribute to mental health vulnerability or resilience later in life. Such elucidated patterns of white matter microstructural disruption might guide future interventions and therapeutic strategies for individuals exposed to ELA. The results of this systematic review are organized by the impacted white matter tracts and the directionality of change in fractional anisotropy. The discussion explores potential mechanisms, highlighting possible reasons why certain pathways are affected whilst others are not and the potential impact of these microstructural alterations, on behaviour and psychopathology.

## 2. Methods

A systematic literature search of the EMBASE, PubMed and PsycINFO, databases was conducted on the 23^rd^ ^of^ April 2024. The search strategy was prospectively registered with PROSPERO, where comprehensive details and a breakdown of the search methodology are available (*PROSPERO ID:* CRD42024526718). This systematic review was conducted according to the Preferred Reporting Items for Systematic reviews and Meta-Analysis (PRISMA) guidelines (Page et al., 2021). The full search terms can be found in the Supplementary Material.

After de-duplication, the titles and abstracts of 1739 papers were screened, and relevant studies were selected. The inclusion criteria were (1) human, (2) adult (18+ years) studies that (3) investigated white matter microstructural differences between participants, who experienced ELA (4) compared to no or varying degrees of ELA (5) through use of TBSS analysis of DTI data and (6) were written in the English language. Studies were excluded if they (1) reported participants with psychiatric diagnosis, (2) were conducted in groups with drug exposure, (3) pertained to pre-partum adversity, (4) were conducted in cohort with family history of alcohol abuse, or (5) familial depression. With regards to inclusion in this review, papers investigating ELA pertaining to any deviation from the expected environment, including both threat and deprivation forms of adversity were included. However, any papers investigating adversity pertaining to pre-partum periods were not included. The rationale for excluding studies that investigate these effects in populations with psychiatric diagnoses or family histories of alcohol abuse or depression is to minimize confounding factors and focus on identifying changes specific to ELA. Following the screening process, which was blinded and completed by two authors (OM and MC), 1699 records were excluded. 40 full texts were reviewed, and 7 papers were included in the synthesis (Fig. 1). Categorization of ELA or non-ELA groups was based on published and validated thresholds. The following information was extracted: study population characteristics, ELA type, and main findings (Table 1). Due to limited review relevant investigations into the other DTI metrics, only FA findings were included in this review. Detailed characteristics of included studies are reported in Table 1 and a quality scoring questionnaire is available in the Supplementary Material.

**Figure 1:**
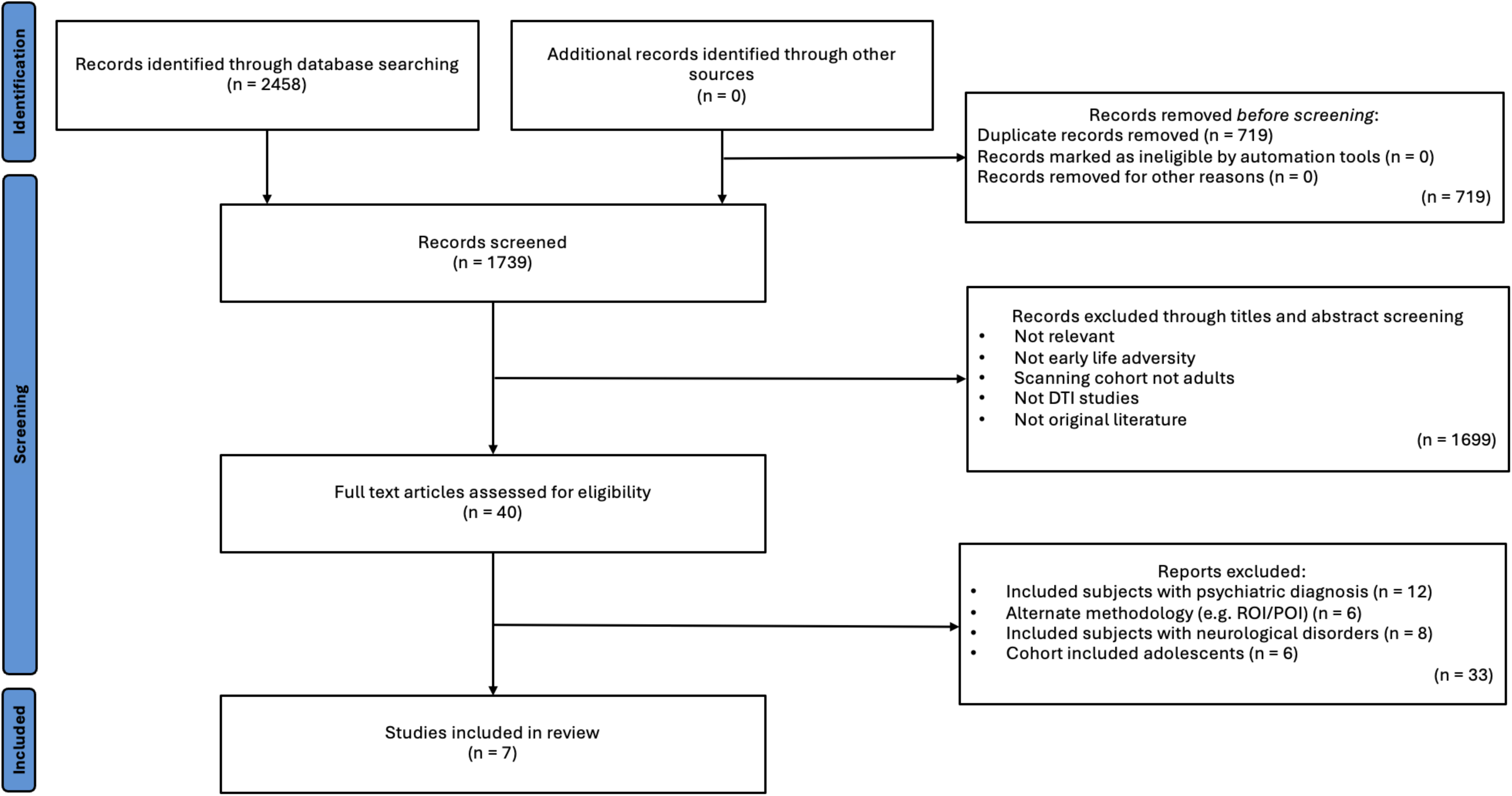
Flow diagram of study selection.

**Table 1:**
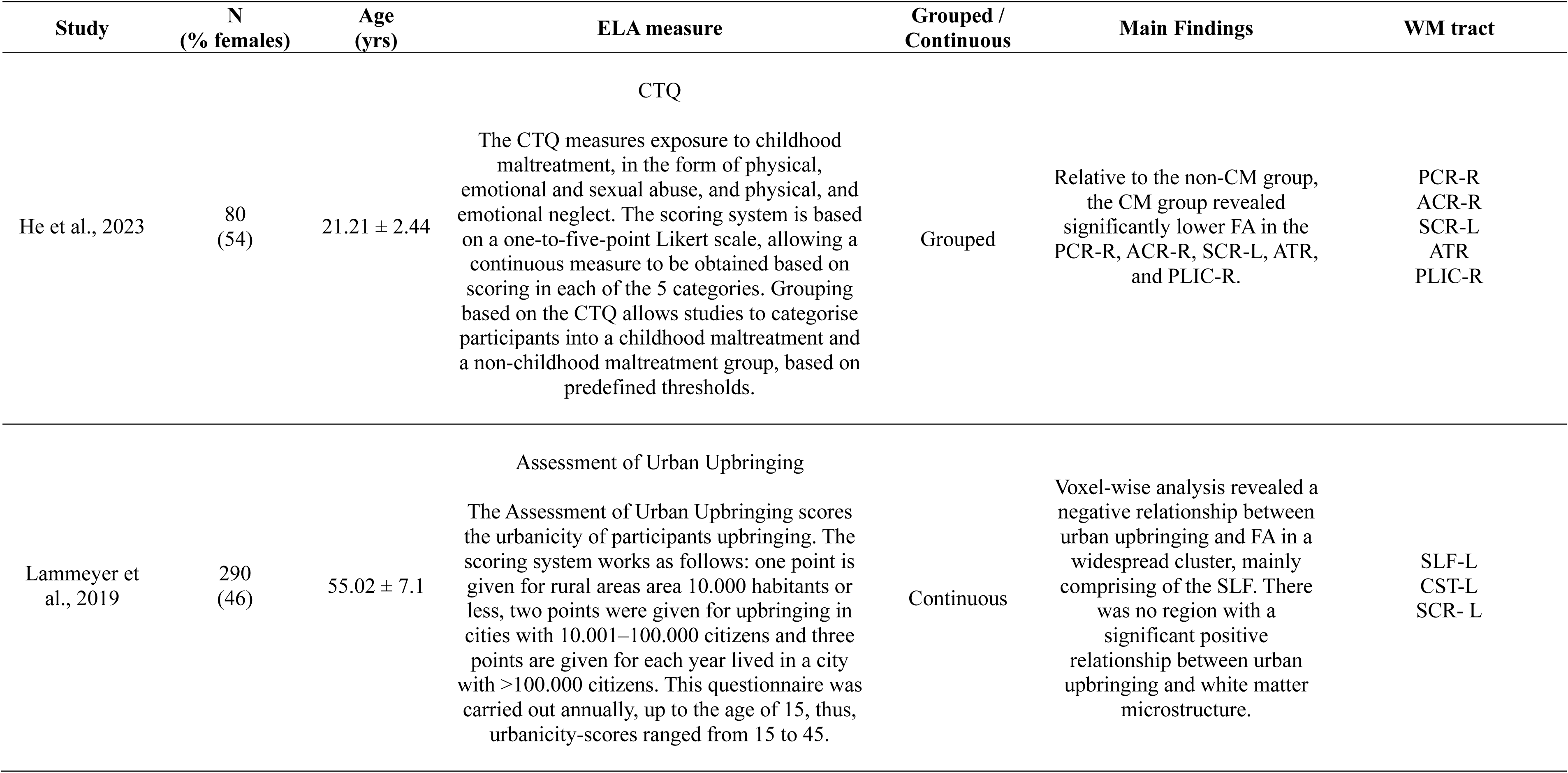
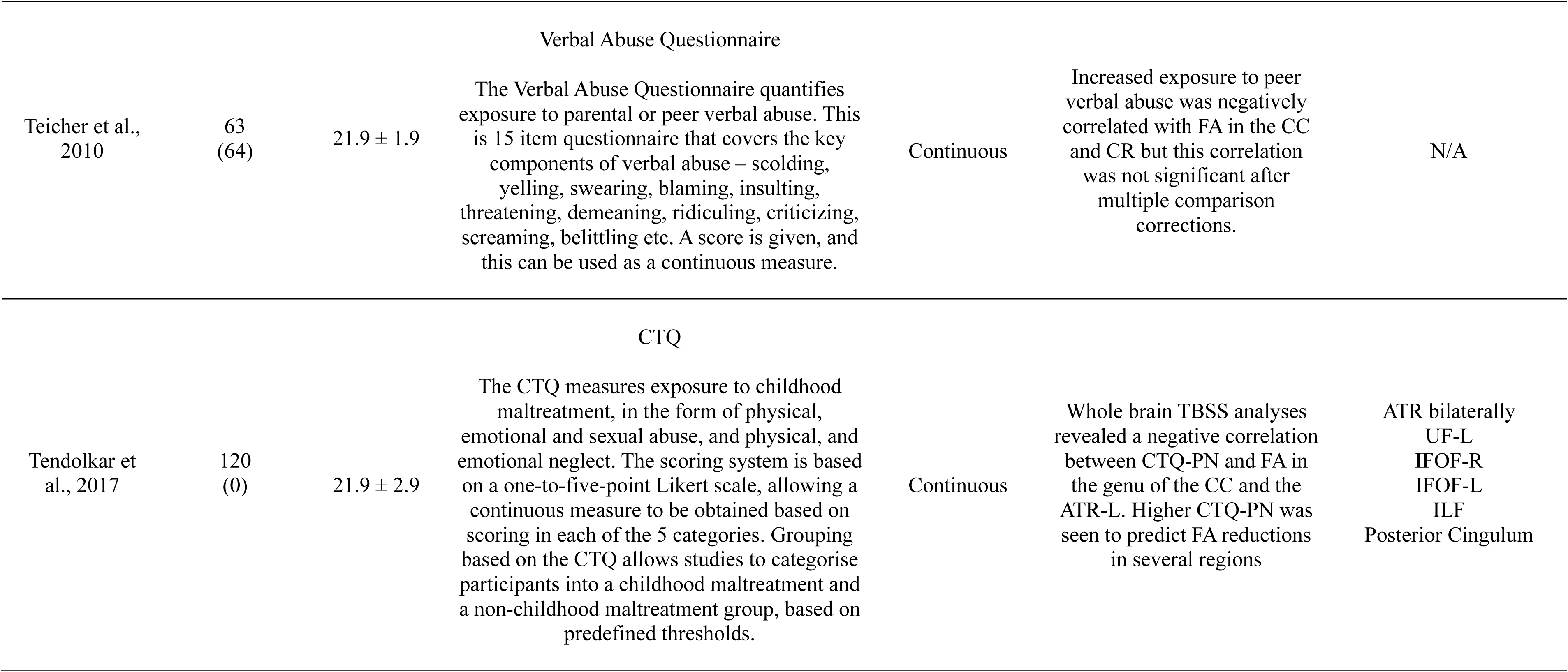
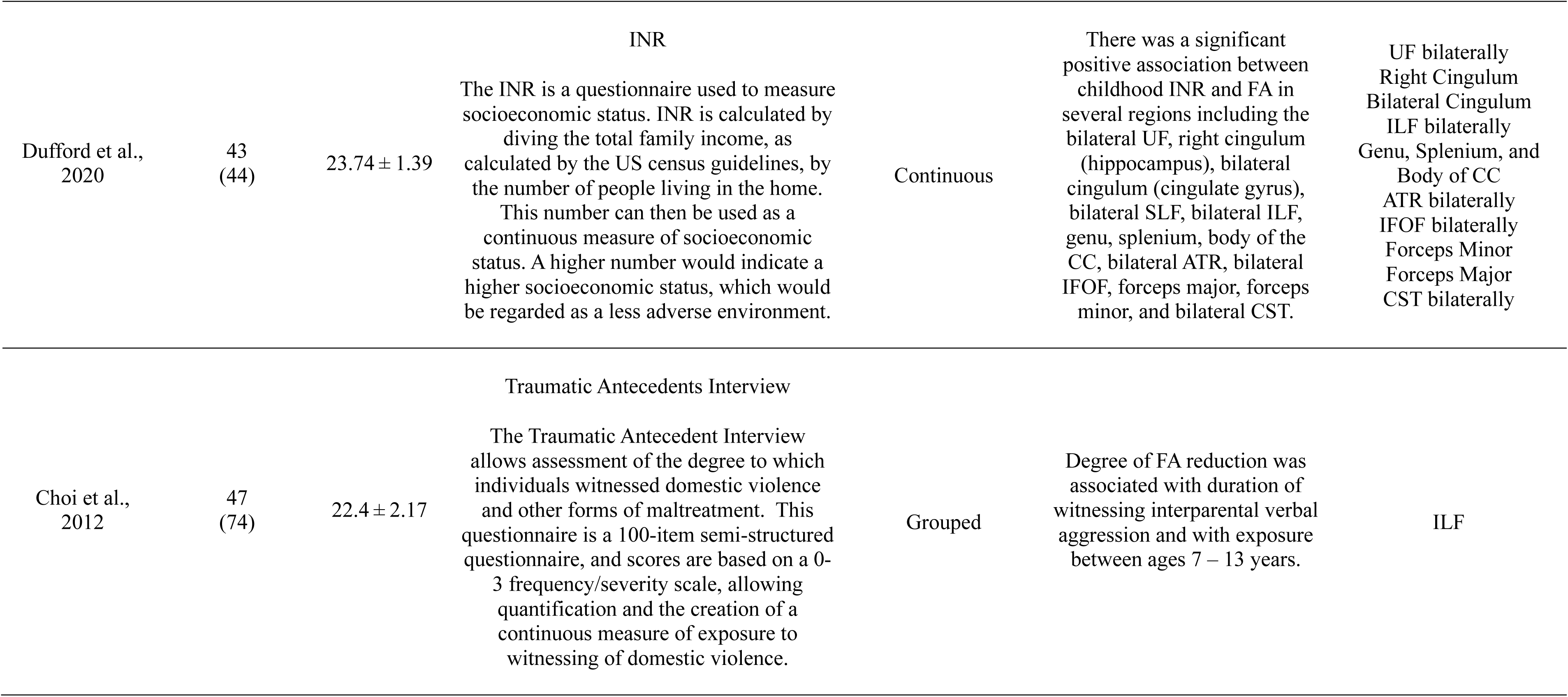
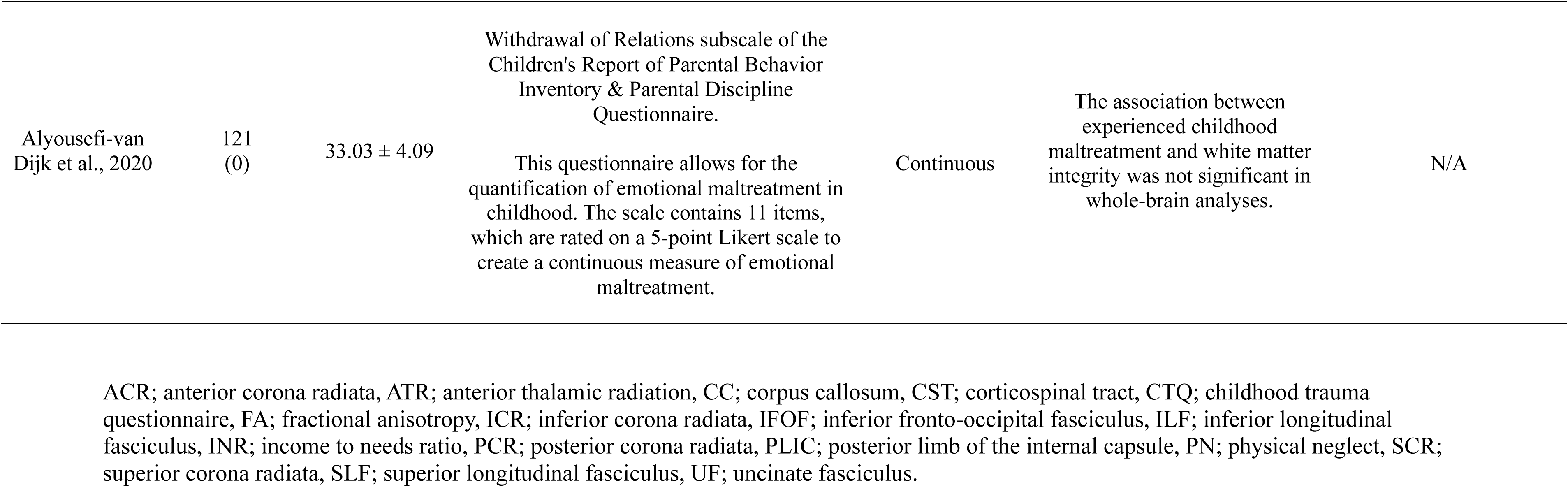
The characteristics of the seven early life adversity studies included in the narrative synthesis.

| Study | N<br>(% females) | Age<br>(yrs) | ELA measure | Grouped /<br>Continuous | Main Findings | WM tract |
| --- | --- | --- | --- | --- | --- | --- |
| CTQ |  |  |  |  |  |  |
| He et al., 2023 | 80<br>(54) | 21.21 ± 2.44 | The CTQ measures exposure to childhood maltreatment, in the form of physical, emotional and sexual abuse, and physical, and emotional neglect. The scoring system is based on a one-to-five-point Likert scale, allowing a continuous measure to be obtained based on scoring in each of the 5 categories. Grouping based on the CTQ allows studies to categorise participants into a childhood maltreatment and a non-childhood maltreatment group, based on predefined thresholds. | Grouped | Relative to the non-CM group, the CM group revealed significantly lower FA in the PCR-R, ACR-R, SCR-L, ATR, and PLIC-R. | PCR-R<br>ACR-R<br>SCR-L<br>ATR<br>PLIC-R |
| Assessment of Urban Upbringing |  |  |  |  |  |  |
| Lammeyer et al., 2019 | 290<br>(46) | 55.02 ± 7.1 | The Assessment of Urban Upbringing scores the urbanicity of participants upbringing. The scoring system works as follows: one point is given for rural areas area 10.000 habitants or less, two points were given for upbringing in cities with 10.001–100.000 citizens and three points are given for each year lived in a city with >100.000 citizens. This questionnaire was carried out annually, up to the age of 15, thus, urbanicity-scores ranged from 15 to 45. | Continuous | Voxel-wise analysis revealed a negative relationship between urban upbringing and FA in a widespread cluster, mainly comprising of the SLF. There was no region with a significant positive relationship between urban upbringing and white matter microstructure. | SLF-L<br>CST-L<br>SCR- L |
| Verbal Abuse Questionnaire |  |  |  |  |  |  |
| Teicher et al.,<br>2010 | 63<br>(64) | 21.9 ± 1.9 | The Verbal Abuse Questionnaire quantifies exposure to parental or peer verbal abuse. This is 15 item questionnaire that covers the key components of verbal abuse – scolding, yelling, swearing, blaming, insulting, threatening, demeaning, ridiculing, criticizing, screaming, belittling etc. A score is given, and this can be used as a continuous measure. | Continuous | Increased exposure to peer verbal abuse was negatively correlated with FA in the CC and CR but this correlation was not significant after multiple comparison corrections. | N/A |
| CTQ |  |  |  |  |  |  |
| Tendolkar et al., 2017 | 120<br>(0) | 21.9 ± 2.9 | The CTQ measures exposure to childhood maltreatment, in the form of physical, emotional and sexual abuse, and physical, and emotional neglect. The scoring system is based on a one-to-five-point Likert scale, allowing a continuous measure to be obtained based on scoring in each of the 5 categories. Grouping based on the CTQ allows studies to categorise participants into a childhood maltreatment and a non-childhood maltreatment group, based on predefined thresholds. | Continuous | Whole brain TBSS analyses revealed a negative correlation between CTQ-PN and FA in the genu of the CC and the ATR-L. Higher CTQ-PN was seen to predict FA reductions in several regions | ATR bilaterally<br>UF-L<br>IFOF-R<br>IFOF-L<br>ILF<br>Posterior Cingulum |
| Dufford et al.,<br>2020 | 43<br>(44) | 23.74 ± 1.39 | INR<br>The INR is a questionnaire used to measure socioeconomic status. INR is calculated by dividing the total family income, as calculated by the US census guidelines, by the number of people living in the home. This number can then be used as a continuous measure of socioeconomic status. A higher number would indicate a higher socioeconomic status, which would be regarded as a less adverse environment. | Continuous | There was a significant positive association between childhood INR and FA in several regions including the bilateral UF, right cingulum (hippocampus), bilateral cingulum (cingulate gyrus), bilateral SLF, bilateral ILF, genu, splenium, body of the CC, bilateral ATR, bilateral IFOF, forceps major, forceps minor, and bilateral CST. | UF bilaterally<br>Right Cingulum<br>Bilateral Cingulum<br>ILF bilaterally<br>Genu, Splenium, and<br>Body of CC<br>ATR bilaterally<br>IFOF bilaterally<br>Forceps Minor<br>Forceps Major<br>CST bilaterally |
| Choi et al.,<br>2012 | 47<br>(74) | 22.4 ± 2.17 | Traumatic Antecedents Interview<br>The Traumatic Antecedent Interview allows assessment of the degree to which individuals witnessed domestic violence and other forms of maltreatment. This questionnaire is a 100-item semi-structured questionnaire, and scores are based on a 0-3 frequency/severity scale, allowing quantification and the creation of a continuous measure of exposure to witnessing of domestic violence. | Grouped | Degree of FA reduction was associated with duration of witnessing interparental verbal aggression and with exposure between ages 7 – 13 years. | ILF |
| Alyousefi-van<br>Dijk et al., 2020 | 121<br>(0) | 33.03 ± 4.09 | Withdrawal of Relations subscale of the Children's Report of Parental Behavior Inventory & Parental Discipline Questionnaire.<br>This questionnaire allows for the quantification of emotional maltreatment in childhood. The scale contains 11 items, which are rated on a 5-point Likert scale to create a continuous measure of emotional maltreatment. | Continuous | The association between experienced childhood maltreatment and white matter integrity was not significant in whole-brain analyses. | N/A |

ACR; anterior corona radiata, ATR; anterior thalamic radiation, CC; corpus callosum, CST; corticospinal tract, CTQ; childhood trauma questionnaire, FA; fractional anisotropy, ICR; inferior corona radiata, IFOF; inferior fronto-occipital fasciculus, ILF; inferior longitudinal fasciculus, INR; income to needs ratio, PCR; posterior corona radiata, PLIC; posterior limb of the internal capsule, PN; physical neglect, SCR; superior corona radiata, SLF; superior longitudinal fasciculus, UF; uncinate fasciculus.

## 3. Results

### 3.1. Study selection

A systematic search of Embase, PubMed and PsycINFO retrieved abstracts of 2,458 English papers on 23 April 2024, which were reduced to seven TBSS studies to compare FA within adults with varying degrees of ELAs or adults exposed to ELA compared to controls (Table 1), involving 764 subjects.

ELA is a general term that encompasses both threat and deprivation realms of adversity. Out of the seven papers included in this review, six investigated ELA associated with threat and one investigated ELA associated with deprivation. Of the seven papers included, four investigated childhood maltreatment, through the childhood trauma questionnaire (He et al., 2023; Tendolkar et al., 2018), verbal abuse questionnaire (Teicher et al., 2004), and withdrawal of relations subscale of the Children’s Report of Parental Behavior Inventory (Alyousefi-van Dijk et al., 2021). One paper looked at an assessment of urban upbringing (Lammeyer et al., 2019), one assessed socio-economic status through an income to needs ratio (Dufford et al., 2020), and one reported adversity as exposure interparental domestic violence (Choi et al., 2012). Each paper’s investigation and findings are detailed in Table 1.

Below, we first summarize the results of a narrative synthesis, according to topographical organization (Schotten, 2012). These can be categorized into projection pathways, which transmit sensory-motor information; association pathways, which integrate information from different brain regions within the same hemisphere; and commissural pathways, which facilitate information transfer between the two hemispheres (Schotten, 2012). Despite not having homogeneity regarding tract, the directionality of FA change was consistent. Of the seven papers reviewed, two found no significant difference in FA, when investigating exposure to verbal abuse and childhood maltreatment (Alyousefi-van Dijk et al., 2021; Teicher et al., 2004). All other papers had significant findings and observed that with increasing adversity, FA decreased in various white matter tracts.

### 3.2 Projection fibers

#### 3.2.1 Corticospinal tract

Two papers implicated the corticospinal tract (CST) in their findings, one on the left (736 voxels, MNI x, y, z: 117, 1141, 95) (Lammeyer et al., 2019) and one bilaterally (Dufford et al., 2020), with reduced FA correlating with increased urban upbringing (Lammeyer et al., 2019) and with reduced INR respectively (Dufford et al., 2020).

##### 3.2.1.1 Corona Radiata

Part of the CST, the corona radiata was implicated in this systematic review. Two studies found reduced FA values in various parts of the corona radiata including, the right posterior corona radiata (PCR-R) (MNI x, y, z: 111, 93, 101), the right anterior corona radiata (ACR-R) (MNI x, y, z: 117, 1141, 95) and the left superior corona radiata (SCR-L) (736 voxels, MNI x, y, z: 117, 1141, 95). One study identified reduced FA values in the PCR-R and the ACR-R in the childhood maltreatment group, compared to controls, whilst two studies reported findings of a reduction in FA in the SCR-L when exposed to adversity in the form of childhood maltreatment and urban upbringing (He et al., 2023; Lammeyer et al., 2019).

##### 3.2.1.2 Internal Capsule

A large portion of the internal capsule (IC) is composed of the CST, however, one study reported a reduction in FA values specifically in the right posterior limb of the internal capsule (PLIC) in a childhood maltreatment group compared to controls (MNI x, y, z: 105, 120, 76) (He et al., 2023).

#### 3.2.2 Thalamic Radiation

Three papers reported a reduction in FA in the anterior thalamic radiation (ATR). These findings were with regards to childhood maltreatment compared to controls (MNI x, y, z: 95, 122, 77) (He et al., 2023), a negative correlation between FA and childhood maltreatment in the form of physical neglect (Tendolkar et al., 2018) and a positive correlation between FA and income to needs ratio (INR) as an assessment of socioeconomic status (SES) (Dufford et al., 2020).

### 3.3 Association fibers

#### 3.3.1 Superior Longitudinal Fasciculus

A decrease in FA in the superior longitudinal fasciculus (SLF) was reported in one of the seven papers, revealing a negative relationship between urban upbringing and FA in a widespread cluster, mainly comprising of the SLF (736 voxels, MNI x, y, z: 117, 1141, 95) (Lammeyer et al., 2019).

#### 3.3.1 Inferior Longitudinal Fasciculus

The inferior longitudinal fasciculus (ILF) was implicated in three of the seven papers. FA in the ILF was correlated with INR (Dufford et al., 2020), degree of FA reduction in the ILF was associated with duration of witnessing interparental verbal aggression (58 voxels, MNI x, y, z: -2, -79, 11) (Choi et al., 2012) and physical neglect predicted a decrease in FA in the ILF (Tendolkar et al., 2018).

#### 3.3.3 Uncinate Fasciculus

The uncinate fasciculus (UF) was implicated both on the left and right sides, and bilaterally in two of the seven papers. It was found that increased levels of physical neglect, as measured by the CTQ, resulted in lower FA values in the left UF (Tendolkar et al., 2018) and that with decrease in INR a decrease in FA was observed bilaterally in the uncinate (Dufford et al., 2020).

#### 3.3.4 Fronto-Occipital Fasciculus

Two papers implicated a change in the fronto-occipital fasciculus, on the left, right, and bilaterally. One paper found that with increased exposure to physical neglect, FA values in the inferior fronto-occipital fasciculus (IFOF) decreased on both the left and right side independently (Tendolkar et al., 2018). Another paper found a significant positive association with INR and FA bilaterally in the IFOF (Dufford et al., 2020).

#### 3.3.5 Cingulum

Again, the cingulum was implicated in two of the seven papers, with an increase in exposure to physical neglect resulting in a reduction in FA in the posterior cingulum, near the precuneus (Tendolkar et al., 2018). Lower INR was also found to be associated with lower FA in the right and bilateral cingulum (Dufford et al., 2020).

### 3.4 Commissural fibers

#### 3.4.1 Corpus Callosum

One paper reported that lower INR was associated with lower FA values in the genu, splenium and body of the corpus callosum (Dufford et al., 2020).

#### 3.4.2 Forceps Major

The forceps major was implicated in one of the seven papers, with lower FA values observed in association with lower INR (Dufford et al., 2020).

#### 3.4.3 Forceps Minor

Lower FA values in the forceps minor was significantly associated with a decrease in INR (Dufford et al., 2020).

## 4. Discussion

This systematic review reported the results of seven papers, exploring the effects of ELA on white matter microstructure as investigated through TBSS analysis of DTI data. The most robust findings were an association between ELA and a reduced FA in the anterior thalamic radiation (ATR) (Dufford et al., 2020; He et al., 2023; Tendolkar et al., 2018), inferior longitudinal fasciculus (ILF) (Choi et al., 2012; Dufford et al., 2020; Tendolkar et al., 2018), corona radiata (He et al., 2023; Lammeyer et al., 2019), corticospinal tract (CST) (Dufford et al., 2020; Lammeyer et al., 2019), uncinate fasciculus (UF) (Dufford et al., 2020; Tendolkar et al., 2018), inferior fronto-occipital fasciculus (IFOF) (Dufford et al., 2020; Tendolkar et al., 2018), and cingulum bundle (Dufford et al., 2020; Tendolkar et al., 2018).

These findings highlight changes specific to the limbic and visual processing system, which are integral to emotional regulation, memory, and sensory integration (Brodal, 2010; Papez, 1937). The implications of these changes are further explored below. Additionally, potential mechanisms are discussed, such as dysregulation of the HPA axis and mechanisms of myelination disruptions. Understanding these microstructural changes and their underlying mechanisms may provide novel insights into neurobiological impacts of ELA, helping inform future research and intervention.

### 4.1 Early life adversity and the limbic system

Previous research on the effects of ELA, have reported structural and functional hinderances in the frontolimbic circuitry (Chahal et al., 2021; Cohen et al., 2006; Hakamata et al., 2022; Marek et al., 2013; Smith & Pollak, 2020). Part of this circuitry, the ATR, is a tract integral to executive functions and the planning of complex behaviors, including emotional processing (Niida et al., 2018). This review found a reduction in FA in the ATR of adults exposed to ELA (Dufford et al., 2020; He et al., 2023; Tendolkar et al., 2018). ATR microstructural disruption has been implicated in psychiatric disorders, particularly, bipolar disorder, in which reductions in FA values have been observed, in comparison to healthy controls, and associated with a reduction in cognitive function (Magioncalda et al., 2016; Oertel-Knöchel et al., 2014; Versace et al., 2008). A reduction in FA in the ATR has also been correlated with depressive symptoms (Shen et al., 2019), further implicating an emotional regulation function of the ATR.

Another component of the frontolimbic circuitry, the UF, was reported to exhibit an association between ELA and a reduction in FA (Dufford et al., 2020; Tendolkar et al., 2018). The UF, a long-range association fiber which connects the frontal and temporal lobes (Von Der Heide et al., 2013), is considered part of the limbic system and although its function remains somewhat unclear, it is thought to play a role in emotional processing and episodic memory (Von Der Heide et al., 2013). The UF is understood to be particularly susceptible to adverse experiences in adolescence due to its late maturation; one of the few white matter tracts to reach full maturation in the third decade (Lebel et al., 2012). Alterations to the microstructure of the UF are also commonly observed in psychiatric disorders which are characterized by emotional dysregulation (Taylor et al., 2007; Zhang et al., 2012). Further to this, studies outside of the scope of this systematic review substantiate reduced UF FA values in individuals exposed to ELA (Eluvathingal et al., 2006; Hanson et al., 2015; Ho et al., 2017), which might explain behavioral dysregulation in adulthood.

The alteration of the frontolimbic circuitry was further implicated through the reduction of FA in the cingulum (Dufford et al., 2020; Tendolkar et al., 2018), regarded as a core part of the limbic system (Dalgleish, 2004). Previously, FA changes in the cingulum bundle have been observed in populations exposed to high levels of trauma (Fani et al., 2012) and these structural changes are suspected to play a role in the emotional dysregulation observed in post-traumatic stress disorder (Bierer et al., 2015; Sanjuan et al., 2013). Tendolkar et al., observed a reduction in FA in the posterior cingulum in adults exposed to physical neglect in childhood which, through mediation analysis, they correlated to trait anxiety levels in adulthood (Tendolkar et al., 2018). The posterior cingulum innervates the dorsal hippocampus and various DTI studies have found white matter microstructural alterations in this region in psychiatric disorders (Fani et al., 2012; Schermuly et al., 2010; Wessa et al., 2009). The cingulum is also thought to have a function associated with spatial navigation and scene processing (Auger & Maguire, 2013), which might explain why the physical neglect group had microstructural changes here, as understanding threat within a scene would be an important environment adaptation.

Previous research by Choi et al., also observed a significant reduction in FA in children exposed to childhood maltreatment in the form of parental verbal abuse in cingulum (Choi et al., 2009). Similarly to the UF, the cingulum appears particularly susceptible to environmental influence, due to its protracted development (Asato et al., 2010; Lebel & Deoni, 2018; Peters et al., 2014). The disruption of these tracts during development has been shown to negatively impact cognitive (Bathelt et al., 2019) and executive functioning (Peters et al., 2014), and has been correlated with increased anxiety and the development major depressive disorder (Chahal et al., 2022; Chahal et al., 2021; Olson et al., 2015).

Two studies found a reduction FA values in various parts of the corona radiata. (He et al., 2023; Lammeyer et al., 2019). The corona radiata plays a role in attentional control (Niogi et al., 2008), with the anterior corona radiata connecting the thalamus to the anterior cingulate cortex and lateral prefrontal cortex (Wakana et al., 2004), aiding in social regulation and understanding of social nuance (Bertoux et al., 2012; O’Callaghan et al., 2016; Tsujimoto et al., 2011). These observations substantiate prior findings by McCarthy-Jones et al., in a childhood maltreatment group compared to controls (McCarthy-Jones et al., 2018).

These observed changes to the frontolimbic white matter microstructure, through changes to the ATR, UF, cingulum and corona radiata, may elucidate a potential pathway through which ELA influences behaviors controlled by this circuitry. This could include emotional regulation and cognitive control and help us better understand the correlation between ELA and increased risk of psychopathology (Juwariah et al., 2022).

### 4.2 Early life adversity and the visual processing system

Shifting from the limbic system, exposure to ELA appears to lead to reduced FA in the white matter tracts of the visual processing system. The ILF mediates transfer of signals from visual areas to the amygdala and hippocampus, mediating the visual processing of emotionally significant stimulus (Catani et al., 2003). A reduction of FA was reported in association with exposure to childhood abuse in the ILF (Lim et al., 2019). Recent work has reported associations with alterations in microstructure of these visual processing regions and psychiatric symptom development (Harnett et al., 2022; Kravitz et al., 2013), which could be to do with the threat learning that occurs through these pathways (Kravitz et al., 2013). The reduction in FA in the ILF was reported when measuring exposure to witnessing of domestic violence and therefore, this FA change could be explained by a plasticity in white matter tracts that are necessary to respond to visual threat, suggesting an environmentally triggered disturbance in normal pathway development, which could underly the emotional consequences of ELA (Lim et al., 2020).

The same paper by Lim et al., who identified a reduction in FA in the ILF, also observed a reduction in FA in the IFOF, as a result of exposure to ELA (Lim et al., 2019). This is substantiated by the findings of Choi et al., who found that increased witnessing of domestic violence in childhood leads to a reduction in FA in the IFOF in adulthood (Choi et al., 2012). The IFOF links the dorsolateral and premotor prefrontal cortices to the posterior regions of the parietal, temporal, and occipital lobes, as well as the caudal cingulate cortex (Frodl et al., 2012). Functionally, it is involved in awareness and executive function (Schmahmann & Pandya, 2007) and has been found to have a reduced FA in patients with major depressive disorder, which has been linked to emotional visual perception deficits (Kieseppä et al., 2010; Phillips et al., 2003).

The SLF, regarded as part of the visuospatial attention network, is important in the initiation of movement and higher order control of body-centered action (Naidich et al., 2013), as well as being involved in the control of eye movement and attention (Janelle et al., 2022). Microstructural changes in the SLF have been reported in psychiatric disorders such as major depressive disorder and schizophrenia, with veterans without major depressive disorder having higher FA values in the SLF, compared to veterans with major depressive disorder (Matthews et al., 2011). Furthermore, a schizophrenia group was reported to have reduced FA compared to controls (Bopp et al., 2017), which was correlated with verbal working memory scores (Bopp et al., 2017). ELA induced alterations to the SLF might help understanding of how the visual system is impacted by stress mechanisms.

The largest white matter tract, the corpus callosum, connects the right and left cerebral hemispheres. The corpus callosum extends anteriorly into the forceps minor, connecting the frontal lobes, and posteriorly into the forceps major, connecting the occipital lobes (Goldstein et al., 2024). A previous paper by McCarthy et al., suggests that the most replicated findings, in ELA and white matter microstructure research, are observed in corpus callosum (McCarthy-Jones et al., 2018). Further to this, alterations to the microstructure of the corpus callosum have been linked to neuropsychological impairments (Fryer et al., 2008), which are broadly similar to the cognitive changes associated with ELA (Pechtel & Pizzagalli, 2011). Therefore, the neuropsychological impact of ELA could be somewhat explained by the changes in corpus callosal microstructure, however, the association between the function of these tracts and psychopathology remains unclear.

This review (He et al., 2023) further substantiates previous correlations between ELA and reduced FA in the internal capsule and its component tracts (Lim et al., 2020; Monteleone et al., 2019). The function of the internal capsule includes occipital connections with higher order visual cortices and the temporal lobe (Zarei et al., 2007). This reduction in FA in the internal capsule could reflect myelination disruption in tracts that transfer visual information, further suggesting some kind of alteration to threat perception.

### 4.4 Neurobiological mechanisms

Exposure to ELA has been shown to alter the activity of the hypothalamic-pituitary-adrenal (HPA) axis, the body’s main stress response system, with increased activation in response to stress (Anacker et al., 2014). Activity in the HPA is governed by the release of corticotropin-releasing hormone (CRH) and vasopressin (AVP), which cause the secretion of adrenocorticotrophic hormone (ACTH), stimulating the secretion of glucocorticoids (cortisol in humans). Cortisol then binds to and activates glucocorticoid receptors (GR), which act to regulate HPA axis activity through a negative feedback mechanism (Fig. 2) (Anacker et al., 2014). Regulation of the HPA axis is important as previous findings suggest that individual reactivity to stress predicts depression risk (Wichers et al., 2009).

**Figure 2:**
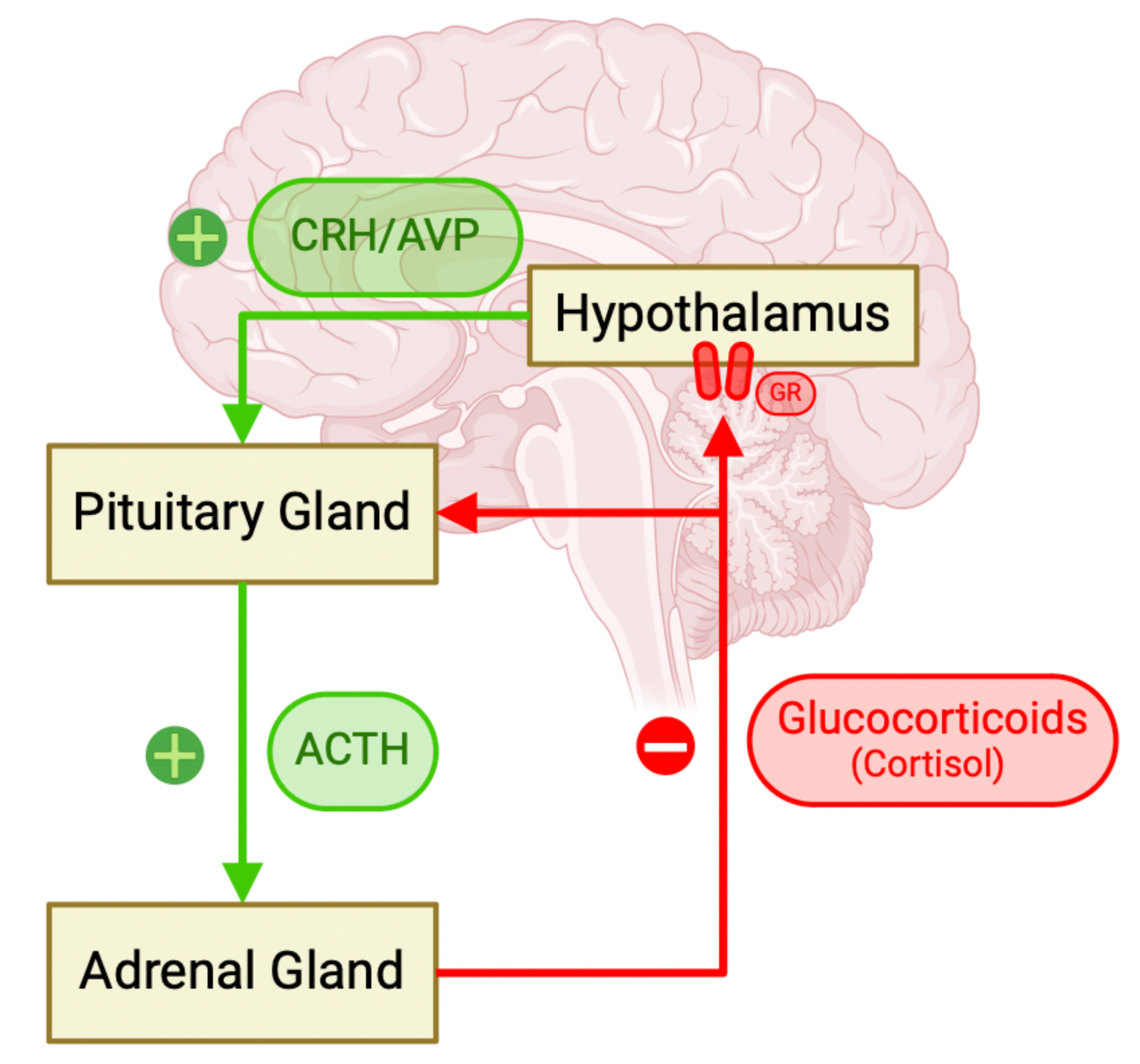
Schematic of hypothalamic-pituitary-adrenal (HPA) axis activity. Image created using www.biorender.com *Legend*. ACTH; adrenocorticotrophic hormone, AVP; vasopressin, CRH; corticotrophin release hormone, GR; glucocorticoid receptor.

It appears that the susceptibility of certain white matter tracts to microstructural alterations as a result of ELA may be due to the timing of maturation and GR mechanisms (Cohodes et al., 2021; Wang et al., 2014). Increased GR expression occurring during adolescence, in response to stress (Isgor et al., 2004), can lead to global cerebral atrophy and reduced FA in white matter (Bourdeau et al., 2002; Chen et al., 2020; van der Meulen et al., 2022; van der Werff et al., 2014). This reduced FA is thought to reflect increased demyelination (Jiang et al., 2017). Demyelination is the process in which the protective myelin sheath surrounding axons is damaged, leading to impaired electrical signal transmission (Love, 2006). Long-term HPA axis activation leads to chronic exposure to cortisol which disrupts oligodendrocyte proliferation and may also leave myelin and lipid membranes vulnerable to oxidative stress (Alonso, 2000; Madrigal et al., 2001; Schiavone et al., 2012). Additionally, cortisol can interact with pro-inflammatory cytokines, disrupting the myelination process (Sapolsky et al., 2000). FA peaking can signify tract maturation; this occurs in certain tracts, such as the ILF, IFOF, UF and cingulum, at early adulthood, opposed to throughout childhood and adolescence. This increased plasticity in early life may lead to increased susceptibility to white matter microstructural changes (Cohodes et al., 2021; Lebel et al., 2012).

### 4.5 Limitations

This study has several limitations. Firstly, this review excluded research within populations with psychiatric diagnoses. This is important to highlight changes resulting from exposure to ELA, whilst removing psychiatric diagnosis as a confounding factor. Despite this, there is a strong link between exposure to ELA and the development of psychiatric disorders (Hughes et al., 2017) meaning excluding these groups may lead to less comprehensive research findings. However, this could provide an interesting subgroup that does not develop psychopathology, perhaps a resilience group. Explorations in this subgroup could provide interesting insights into white matter biomarkers of resilience to psychopathology when exposed to ELA.

One of the strongest protective factors against the development of psychopathology, in the context of ELA, is a stable and supportive caregiving environment (Gee, 2021). Whilst abuse and neglect inherently disrupt caregiving stability, it is an essential covariate in deprivation-based ELA studies. Neither study investigating INR nor urbanicity included this covariate, potentially overlooking a resilience mechanism. This resilience may influence how ELA affects white matter microstructure, therefore, biasing these findings.

Despite being introduced to address VBM-like analysis shortcomings, TBSS has its own limitations. Limitations include anatomical inaccuracies, biases in skeleton projection, limited statistical power, and sensitivity to pathology (Bach et al., 2014). These issues can result in incorrectly identified tracts, partial volume errors, and inaccurate measurements (Figley et al., 2022; Keihaninejad et al., 2012). TBSS’s limited sensitivity and susceptibility to partial volume effects are exacerbated when handling complex white matter structures, often misinterpreting crossing or kissing fibers as reduced FA in one tract. FA values may also be distorted in voxels near tissue boundaries. Despite these challenges, TBSS remains widely used and clear guidelines can address its limitations (Bach et al., 2014). Its popularity allows for reproducibility, and these limitations aren’t always solved by alternative methodologies (Bach et al., 2014).

### 4.6 Future Directions

This review highlights significant white matter microstructural changes in adults exposed to ELA but notes a key gap in research on deprivation-based ELA, despite its social prevalence. Only one of the seven included studies examined deprivation. More research is needed to distinguish brain changes associated specifically with deprivation versus threat-related ELA and to understand the unique white matter alterations from different adversity types, clarifying how ELA influences later psychopathology. Comparing white matter microstructural changes in adults with and without psychiatric diagnoses may reveal resilience factors. Integrating these findings with public health research could help inform policy, underscoring the need for a comprehensive approach to ELA’s varied neurodevelopmental impacts.

## 5. Conclusion

Widespread reductions in white matter microstructure in adults, indicated by a reduced fractional anisotropy, were associated with exposure to ELA, particularly in tracts of the limbic and visual processing networks. These white matter disruptions seem to be a fundamental feature of exposure to ELA and may provide an explanation for the behavioral changes and increased risk of psychopathology observed in adulthood. From these findings, future research can be directed towards the limbic and visual processing systems. Due to the variability of events considered adversity, further research should look to categorize adverse events to allow for more precise findings.

## Declarations

### Ethics Approval

As this study is a systematic review of published literature, no ethical approval was required. All sources referenced in this review were obtained from publicly available, peer-reviewed publications.

### Consent to participate

N/A

### Consent for publication

N/A

### Availability of data and material (data transparency)

N/A

### Code availability

N/A

### Author Contributions

O.M: end-to-end search completion, finding synthesis and interpretation, manuscript writing. M.C: finding synthesis and interpretation, critical revision of manuscript, study supervision. D.R: Study supervision.

