## Supplementary Material for "Early life adversity and White Matter Microstructural Organization - a systematic review"

### <u>Search Strategy</u>

#### EMBASE

23/04/2024 12.13pm BST
1,459 results
June 1998 – April 2024

1. 'child'/exp OR 'child' OR 'childhood'/exp OR 'childhood' OR 'children'/exp OR 'children' OR 'school age'/exp OR 'school age' OR 'early life'/exp OR 'early life' OR 'infant'/exp OR 'infant' OR 'infancy'/exp OR 'infancy' OR 'toddler'/exp OR 'toddler' OR 'youth'/exp OR 'youth' OR 'adolescent'/exp OR 'adolescent' OR 'adolescence'/exp OR 'adolescence' OR 'childhood development' OR 'early childhood development'/exp OR 'early childhood development' OR 'adolescent development'/exp OR 'adolescent development' OR 'adolescent psychology'/exp OR 'adolescent psychology' OR 'developmental psychology'/exp OR 'developmental psychology' OR 'child psychology'/exp OR 'child psychology' OR 'infant development'/exp OR 'infant development' OR 'neonatal development'/exp OR 'neonatal development' OR 'preschool'/exp OR 'preschool'

2. 'adversity' OR 'adverse experience' OR 'adverse childhood experience' OR 'aces' OR 'early-life stress' OR 'els' OR 'maltreatment' OR 'maltreated' OR 'psychological trauma stress' OR 'chronic stress' OR 'environmental stress' OR 'psychological stress' OR 'social stress' OR 'abuse' OR 'abused' OR 'domestic violence' OR 'bullying' OR 'victim' OR 'victimization' OR 'victimisation' OR 'victimized' OR 'victimised' OR 'trauma' OR 'traumatized' OR 'traumatised' OR 'traumatic' OR 'intimate partner violence' OR 'spouse abuse' OR 'sexual abuse' OR 'child abuse' OR 'exposure to violence' OR 'physical abuse' OR 'family conflict' OR 'battered child syndrome' OR 'institutional' OR 'institutional rearing' OR 'institutionalization' OR 'institutionalisation' OR 'institutionalized' OR 'institutionalised' OR 'orphan' OR 'orphanage' OR 'foster care' OR 'adoption' OR 'neglect' OR 'enrichment' OR 'cognitive stimulation' OR 'social stimulation' OR 'residential care institutions' OR 'child neglect' OR 'social deprivation' OR 'abandonment' OR 'psychosocial deprivation' OR 'emotional abuse' OR 'pedophilia' OR 'incest' OR 'rape' OR 'partner abuse' OR 'physical discipline' OR 'socioeconomic' OR 'ses' OR 'poverty' OR 'low income' OR 'low-income' OR 'neighbourhood' OR 'income' OR 'disadvantage' OR 'hardship' OR 'stressful experiences' OR 'maternal depression'

3. 'tbss' OR 'tract based spatial statistics' OR 'whole brain analysis' OR 'diffusion imaging' OR 'diffusion tensor imaging' OR 'white matter microstructure' OR 'white matter properties' OR 'white matter integrity' OR 'fractional anisotropy' OR 'mean diffusivity' OR 'radial diffusivity' OR 'axial diffusivity'

4. #1 AND #2

5. #3 AND #4

PubMed
23/04/2024 10.11am BST
968 results
June 1998 – April 2024

1. ("Child"[All Fields] OR "Childhood"[All Fields] OR "Children"[All Fields] OR "School Age"[All Fields] OR "Early life"[All Fields] OR "Infant"[All Fields] OR "Infancy"[All Fields] OR "Toddler"[All Fields] OR "Youth"[All Fields] OR "Adolescent"[All Fields] OR "Adolescence"[All Fields] OR "Childhood Development"[All Fields] OR "Early Childhood Development"[All Fields] OR "Adolescent Development"[All Fields] OR "Adolescent Psychology"[All Fields] OR "Developmental Psychology"[All Fields] OR "Child Psychology"[All Fields] OR "Infant Development"[All Fields] OR "Neonatal Development"[All Fields] OR "Preschool"[All Fields])

2. ("Adversity"[All Fields] OR "Adverse experience"[All Fields] OR "Adverse childhood experience"[All Fields] OR "ACES"[All Fields] OR "Early-life stress"[All Fields] OR "ELS"[All Fields] OR "Maltreatment"[All Fields] OR "Maltreated"[All Fields] OR "Psychological Stress"[All Fields] OR "Psychological Trauma Stress"[All Fields] OR "Chronic Stress"[All Fields] OR "Environmental Stress"[All Fields] OR "Psychological Stress"[All Fields] OR "Social Stress"[All Fields] OR "Abuse"[All Fields] OR "Abused"[All Fields] OR "Domestic violence"[All Fields] OR "bullying"[All Fields] OR "Victim"[All Fields] OR "Victimization"[All Fields] OR "Victimisation"[All Fields] OR "Victimized"[All Fields] OR "Victimised"[All Fields] OR "Trauma"[All Fields] OR "Traumatized"[All Fields] OR "Traumatised"[All Fields] OR "Traumatic"[All Fields] OR "Intimate Partner Violence"[All Fields] OR "Physical Abuse"[All Fields] OR "Spouse Abuse"[All Fields] OR "Sexual Abuse"[All Fields] OR "Child Abuse"[All Fields] OR "Exposure to Violence"[All Fields] OR "Rape"[All Fields] OR "Physical Abuse"[All Fields] OR "Family Conflict"[All Fields] OR "Battered Child Syndrome"[All Fields] OR "Institutional"[All Fields] OR "Institutional rearing"[All Fields] OR "Institutionalization"[All Fields] OR "Institutionalisation"[All Fields] OR "Institutionalized"[All Fields] OR "Institutionalised"[All Fields] OR "Orphan"[All Fields] OR "Orphanage"[All Fields] OR "foster care"[All Fields] OR "adoption"[All Fields] OR "Neglect"[All Fields] OR "Enrichment"[All Fields] OR "Cognitive Stimulation"[All Fields] OR "Social Stimulation"[All Fields] OR "Residential Care Institutions"[All Fields] OR "Child Neglect"[All Fields] OR "Social Deprivation"[All Fields] OR "Abandonment"[All Fields] OR "Psychosocial Deprivation"[All Fields] OR "Emotional Abuse"[All Fields] OR "Pedophilia"[All Fields] OR "Incest"[All Fields] OR "Rape"[All Fields] OR "Partner Abuse"[All Fields] OR "Physical Discipline"[All Fields] OR "socioeconomic"[All Fields] OR "SES"[All Fields] OR "poverty"[All Fields] OR "low-income"[All Fields] OR "low-income"[All Fields] OR "neighbourhood"[All Fields] OR "income"[All Fields] OR "disadvantage"[All Fields] OR "hardship"[All Fields] OR "stressful experiences"[All Fields] OR "maternal depression"[All Fields])

3. ("TBSS"[All Fields] OR "tract based spatial statistics"[All Fields] OR "whole brain analysis"[All Fields] OR "diffusion imaging"[All Fields] OR "diffusion tensor imaging"[All Fields] OR "white matter microstructure"[All Fields] OR "white matter properties"[All Fields] OR "white matter integrity"[All Fields] OR "fractional anisotropy"[All Fields] OR "mean diffusivity"[All Fields] OR "radial diffusivity"[All Fields] OR "axial diffusivity"[All Fields])

4. #1 AND #2

5. #3 AND #4

PsycINFO
23/04/2024 13.00pm BST
31 results
June 2013 – Feb 2024

1. 'child'/exp OR 'child' OR 'childhood'/exp OR 'childhood' OR 'children'/exp OR 'children' OR 'school age'/exp OR 'school age' OR 'early life'/exp OR 'early life' OR 'infant'/exp OR 'infant' OR 'infancy'/exp OR 'infancy' OR 'toddler'/exp OR 'toddler' OR 'youth'/exp OR 'youth' OR 'adolescent'/exp OR 'adolescent' OR 'adolescence'/exp OR 'adolescence' OR 'childhood development' OR 'early childhood development'/exp OR 'early childhood development' OR 'adolescent development'/exp OR 'adolescent development' OR 'adolescent psychology'/exp OR 'adolescent psychology' OR 'developmental psychology'/exp OR 'developmental psychology' OR 'child psychology'/exp OR 'child psychology' OR 'infant development'/exp OR 'infant development' OR 'neonatal development'/exp OR 'neonatal development' OR 'preschool'/exp OR 'preschool'

2. 'adversity' OR 'adverse experience' OR 'adverse childhood experience' OR 'aces' OR 'early-life stress' OR 'els' OR 'maltreatment' OR 'maltreated' OR 'psychological trauma stress' OR 'chronic stress' OR 'environmental stress' OR 'psychological stress' OR 'social stress' OR 'abuse' OR 'abused' OR 'domestic violence' OR 'bullying' OR 'victim' OR 'victimization' OR 'victimisation' OR 'victimized' OR 'victimised' OR 'trauma' OR 'traumatized' OR 'traumatised' OR 'traumatic' OR 'intimate partner violence' OR 'spouse abuse' OR 'sexual abuse' OR 'child abuse' OR 'exposure to violence' OR 'physical abuse' OR 'family conflict' OR 'battered child syndrome' OR 'institutional' OR 'institutional rearing' OR 'institutionalization' OR 'institutionalisation' OR 'institutionalized' OR 'institutionalised' OR 'orphan' OR 'orphanage' OR 'foster care' OR 'adoption' OR 'neglect' OR 'enrichment' OR 'cognitive stimulation' OR 'social stimulation' OR 'residential care institutions' OR 'child neglect' OR 'social deprivation' OR 'abandonment' OR 'psychosocial deprivation' OR 'emotional abuse' OR 'pedophilia' OR 'incest' OR 'rape' OR 'partner abuse' OR 'physical discipline' OR 'socioeconomic' OR 'ses' OR 'poverty' OR 'low income' OR 'low-income' OR 'neighbourhood' OR 'income' OR 'disadvantage' OR 'hardship' OR 'stressful experiences' OR 'maternal depression'

3. 'tbss' OR 'tract based spatial statistics' OR 'whole brain analysis' OR 'diffusion imaging' OR 'diffusion tensor imaging' OR 'white matter microstructure' OR 'white matter properties' OR 'white matter integrity' OR 'fractional anisotropy' OR 'mean diffusivity' OR 'radial diffusivity' OR 'axial diffusivity'

4. #1 AND #2

5. #3 AND #4

#### Table 1. White Matter Tracts implicated, their anatomy and function.

| Tract | Anatomy | Function |
|---|---|---|
| **Corticospinal Tract** | The corticospinal tract is a projection fibre that connects the spinal cord to the cortex. Originating in the motor cortex, the corticospinal tract travels through the internal capsule and the cerebral peduncles, down through the brainstem, terminating in the anterior horn of the spinal cord where it synapses with motor neurons (Jang, 2014). | The primary pathway for voluntary motor control, enabling movement of the body, especially fine motor control of the limbs (Rong et al., 2014). |
| **Corona Radiata** | The corona radiata is a projection fibre, composed of a collection of white matter tracts connecting the cortex to subcortical structures and terminates at the superior border of the lentiform nucleus. The corona radiata then converges into the internal capsule, caudally (Costa et al., 2018). | The corona radiata acts as a major conduit for ascending sensory information to the cortex and descending motor signals from the cortex, facilitating communication between cortical and subcortical regions (Javed et al., 2024). |
| **Internal Capsule** | The internal capsule is continuation of the corona radiata inferiorly, another projection fibre, that connects the cerebral hemispheres with subcortical structures, the brainstem and spinal cord and is made up of both motor and sensory fibres. It traverses the basal ganglia, dividing the caudate nucleus and thalamus from the putamen and globus pallidus (Emos et al., 2024). | Plays a role in both motor control and sensory processing but transmitting motor and sensory information between the cerebral cortex and spinal cord or brainstem (Emos et al., 2024). |
| **Thalamic Radiation** | The thalamic radiations are projection fibres, that connect the thalamus to the cerebral cortex. There are four distinct thalamic radiations; the inferior, superior, anterior, and posterior thalamic radiations (George & Das, 2024). | The various thalamic radiations act to relay sensory or motor information from the thalamus to distinct areas of the cerebral cortex. The thalamus is often referred to as the relay centre of the brain, sending information to appropriate regions of the cortex, via the thalamic radiations (George & Das, 2024). |

| | | |
|---|---|---|
| **Superior Longitudinal Fasciculus** | The superior longitudinal fasciculus is an association fibre that connects the frontal lobe and the other lobes (occipital, parietal, and temporal) of the ipsilateral hemisphere (Janelle et al., 2022). | The superior longitudinal fasciculus has functions related to language, attention, and spatial processing, as well as the integration of sensory-motor information (Janelle et al., 2022). |
| **Inferior Longitudinal Fasciculus** | The inferior longitudinal fasciculus, an association pathway, extends from the occipital pole to the temporal pole, passing through the fusiform gyrus and transfers visual information from the visual areas to the amygdala and hippocampus. | Primary function is the processing of visual information, including working memory for the short-term maintenance of information (Catani et al., 2003; Patel et al., 2024). |
| **Uncinate Fasciculus** | The uncinate fasciculus is an association pathway, that connects the orbitofrontal cortex to the anterior part of the temporal lobe, passing by the Sylvian fissure inferiorly. | Although the function of the uncinate fasciculus remains somewhat unclear, it is thought to play a role in emotional processing and episodic memory (Von Der Heide et al., 2013). |
| **Fronto-Occipital Fasciculus** | The fronto-occipital fasciculus, also an association fibre, connects the dorsolateral and premotor prefrontal cortices to the posterior regions of the parietal, temporal, and occipital lobes, as well as the caudal cingulate cortex (Frodl et al., 2012). | The function of the fronto-occipital fasciculus appears to be related to awareness and executive function (Schmahmann et al., 2007), as well as sematic and visual processing (Wu et al., 2020). |
| **Cingulum** | The cingulum is a c-shaped association fibre bundle that lies within the cingulate gyrus and curves around the corpus callosum, acting to connect the frontal, parietal and temporal lobes., as well as connecting subcortical nuclei to the cingulate gyrus (Bubb et al., 2018). | Involved in emotional regulation, memory, and cognitive control, the cingulum connects the limbic areas with the cortical areas. |

| | | |
|---|---|---|
| **Corpus Callosum** | The corpus callosum, a commissural fibre, connects the left and right hemispheres of the brain (Goldstein et al., 2024). | Primarily, the function of the corpus callosum is to integrate and transfer information from each of the cerebral hemispheres, to facilitate the processing of sensory, motor and cognitive stimulus. |
| **Forceps Major** | The forceps major is the most posterior part of the corpus callosum. Many of the fibres of the splenium of the corpus callosum travel posteriorly into the occipital lobe to form the forceps major, a similar, commissural fibre (Goldstein et al., 2024). | The fibres of the forceps minor form a connection between regions of the occipital lobes, therefore, playing a role in visual information processing and integration of this information across hemispheres. |
| **Forceps Minor** | The forceps minor is the most anterior part of the corpus callosum. Many fibres from the genu of the corpus callosum, travel anteriorly into the frontal lobes. These fibres on the two hemispheres of the brain form the forceps minor, a fork-like, commissural fibre (Goldstein et al., 2024). | The fibres of the forceps major form a connection between regions of the frontal cortices. This permits interhemispheric communication for the frontal lobes, allowing integration of information associated with executive function and cognition, across the two hemispheres. |

### Quality Assessment Checklist

Imaging Methodology Quality Assessment Checklist adapted from (Disorganization of white matter architecture in major depressive disorder: a meta-analysis of diffusion tensor imaging with tract-based spatial statistics. - Guangxiang Chen, Xinyu Hu, Lei Li, Xiaoqi Huang, Su Lui, Weihong Kuang, Hua Ai, Feng Bi, Zhongwei Gu & Qiyong Gong)

Supplementary Table S1: Imaging Methodology Quality Assessment Checklist
(When criteria were partially met, 0.5 points were assigned)

| **Category 1: Subjects** | **Score** (0/0.5/1) |
|---|---|
| 1. Patients were recruited prospectively and demographic data was reported | |
| 2. Important variables (e.g. age, gender, race) were compared between the groups, either by stratification or statistically | |
| 3. Sample size per group > 10 | |
| **Category 2: Methods for image acquisition and analysis** | |
| 4. Magnet strength at least 1.5T | |
| 5. MRI slice-thickness≤3 mm | |
| 6. Whole brain analysis was automated with no a-priori regional selection | |
| 7. Coordinates reported in a standard space | |
| 8. The imaging technique used was clearly described so that it could be reproduced | |
| 9. Measurements were clearly described so that they could be reproduced | |
| **Category 3: Results and conclusions** | |
| 10. Statistical parameters (p-value, cluster size, cluster coordinates) for significant differences were provided | |
| 11. Conclusions were consistent with the results obtained and the limitations were discussed | |
| | **TOTAL** /11 |

Supplementary Table S2: Imaging Methodology Quality Assessment Checklist for paper 1.

"Characteristics of white matter structural connectivity in healthy adults with childhood maltreatment" - Jiayue He, Xue Zhong, Chang Cheng, Daifeng Dong, Bei Zhang, Xiang Wang, Shuqiao Yaoa.

| **Category 1: Subjects** | **Score** (0/0.5/1) |
|---|---|
| 1. Patients were recruited prospectively and demographic data was reported (1) | |
| 2. Important variables (e.g. age, gender, race) were compared between the groups, either by stratification or statistically (1) | |
| 3. Sample size per group > 10 (1) | |
| **Category 2: Methods for image acquisition and analysis** | |
| 4. Magnet strength at least 1.5T (1) | |
| 5. MRI slice-thickness≤3 mm (1) | |
| 6. Whole brain analysis was automated with no a-priori regional selection (1) | |
| 7. Coordinates reported in a standard space (0) | |
| 8. The imaging technique used was clearly described so that it could be reproduced (1) | |
| 9. Measurements were clearly described so that they could be reproduced (1) | |
| **Category 3: Results and conclusions** | |
| 10. Statistical parameters (p-value, cluster size, cluster coordinates) for significant differences were provided (1) | |
| 11. Conclusions were consistent with the results obtained and the limitations were discussed (1) | |
| | **TOTAL** 9 /11 |

#### Supplementary Table S2: Imaging Methodology Quality Assessment Checklist for paper 2.

"Evidence of brain network aberration in healthy subjects with urban upbringing – A multimodal DTI and VBM study" – Sophia Lammeyer, Bruno Dietsche, Udo Dannlowski, Tilo Kircher, Axel Krug.

| **Category 1: Subjects** | **Score** (0/0.5/1) |
|---|---|
| 1. Patients were recruited prospectively and demographic data was reported (0.5) | |
| 2. Important variables (e.g. age, gender, race) were compared between the groups, either by stratification or statistically (0) | |
| 3. Sample size per group > 10 (1) | |
| **Category 2: Methods for image acquisition and analysis** | |
| 4. Magnet strength at least 1.5T (1) | |
| 5. MRI slice-thickness≤3 mm (1) | |
| 6. Whole brain analysis was automated with no a-priori regional selection (1) | |
| 7. Coordinates reported in a standard space (1) | |
| 8. The imaging technique used was clearly described so that it could be reproduced (1) | |
| 9. Measurements were clearly described so that they could be reproduced (1) | |
| **Category 3: Results and conclusions** | |
| 10. Statistical parameters (p-value, cluster size, cluster coordinates) for significant differences were provided (1) | |
| 11. Conclusions were consistent with the results obtained and the limitations were discussed (1) | |
| | **TOTAL** 9.5/11 |

Supplementary Table S2: Imaging Methodology Quality Assessment Checklist for paper 3.

"Hurtful Words: Association of Exposure to Peer Verbal Abuse with Elevated Psychiatric Symptom Scores and Corpus Callosum Abnormalities" - Martin H Teicher, Jacqueline A Samson, Yi-Shin Sheu, Ann Polcari, Cynthia E McGreenery.

| **Category 1: Subjects** | **Score** (0/0.5/1) |
|---|---|
| 1. Patients were recruited prospectively and demographic data was reported (0) | |
| 2. Important variables (e.g. age, gender, race) were compared between the groups, either by stratification or statistically (0) | |
| 3. Sample size per group > 10 (1) | |
| **Category 2: Methods for image acquisition and analysis** | |
| 4. Magnet strength at least 1.5T (1) | |
| 5. MRI slice-thickness≤3 mm (0) | |
| 6. Whole brain analysis was automated with no a-priori regional selection (1) | |
| 7. Coordinates reported in a standard space (1) | |
| 8. The imaging technique used was clearly described so that it could be reproduced (1) | |
| 9. Measurements were clearly described so that they could be reproduced (1) | |
| **Category 3: Results and conclusions** | |
| 10. Statistical parameters (p-value, cluster size, cluster coordinates) for significant differences were provided (0.5) | |
| 11. Conclusions were consistent with the results obtained and the limitations were discussed (1) | |
| | **TOTAL** 7.5 /11 |

Supplementary Table S2: Imaging Methodology Quality Assessment Checklist for paper 4.

"Physical neglect during childhood alters white matter connectivity in healthy young males" - Indira Tendolkar, Johan Mårtensson, Simone Kühn, Floris Klumpers, Guillén Fernández.

| **Category 1: Subjects** | **Score** (0/0.5/1) |
|---|---|
| 1. Patients were recruited prospectively and demographic data was reported (1) | |
| 2. Important variables (e.g. age, gender, race) were compared between the groups, either by stratification or statistically (0) | |
| 3. Sample size per group > 10 (1) | |
| **Category 2: Methods for image acquisition and analysis** | |
| 4. Magnet strength at least 1.5T (1) | |
| 5. MRI slice-thickness≤3 mm (1) | |
| 6. Whole brain analysis was automated with no a-priori regional selection (1) | |
| 7. Coordinates reported in a standard space (0) | |
| 8. The imaging technique used was clearly described so that it could be reproduced (1) | |
| 9. Measurements were clearly described so that they could be reproduced (1) | |
| **Category 3: Results and conclusions** | |
| 10. Statistical parameters (p-value, cluster size, cluster coordinates) for significant differences were provided (0) | |
| 11. Conclusions were consistent with the results obtained and the limitations were discussed (1) | |
| | **TOTAL** 8 /11 |

Supplementary Table S2: Imaging Methodology Quality Assessment Checklist for paper 5.

"Prospective associations, longitudinal patterns of childhood socioeconomic status, and white matter organization in adulthood" - Alexander J Dufford, Gary W Evans, Julia Dmitrieva, James E Swain, Israel Liberzon, Pilyoung Kim.

| **Category 1: Subjects** | **Score** (0/0.5/1) |
|---|---|
| 1. Patients were recruited prospectively and demographic data was reported (1) | |
| 2. Important variables (e.g. age, gender, race) were compared between the groups, either by stratification or statistically (1) | |
| 3. Sample size per group > 10 (1) | |
| **Category 2: Methods for image acquisition and analysis** | |
| 4. Magnet strength at least 1.5T (1) | |
| 5. MRI slice-thickness≤3 mm (1) | |
| 6. Whole brain analysis was automated with no a-priori regional selection (1) | |
| 7. Coordinates reported in a standard space (0) | |
| 8. The imaging technique used was clearly described so that it could be reproduced (1) | |
| 9. Measurements were clearly described so that they could be reproduced (1) | |
| **Category 3: Results and conclusions** | |
| 10. Statistical parameters (p-value, cluster size, cluster coordinates) for significant differences were provided (0.5) | |
| 11. Conclusions were consistent with the results obtained and the limitations were discussed (1) | |
| | **TOTAL** 9.5 /11 |

#### Supplementary Table S2: Imaging Methodology Quality Assessment Checklist for paper 6.

"Reduced Fractional Anisotropy in the Visual Limbic Pathway of Young Adults Witnessing Domestic Violence in Childhood" - Jeewook Choi, Bumseok Jeong, Ann Polcari, Michael L Rohan, Martin H Teicher.

| **Category 1: Subjects** | **Score** (0/0.5/1) |
|---|---|
| 1. Patients were recruited prospectively and demographic data was reported (1) | |
| 2. Important variables (e.g. age, gender, race) were compared between the groups, either by stratification or statistically (1) | |
| 3. Sample size per group > 10 (1) | |
| **Category 2: Methods for image acquisition and analysis** | |
| 4. Magnet strength at least 1.5T (1) | |
| 5. MRI slice-thickness≤3 mm (0) | |
| 6. Whole brain analysis was automated with no a-priori regional selection (1) | |
| 7. Coordinates reported in a standard space (1) | |
| 8. The imaging technique used was clearly described so that it could be reproduced (1) | |
| 9. Measurements were clearly described so that they could be reproduced (1) | |
| **Category 3: Results and conclusions** | |
| 10. Statistical parameters (p-value, cluster size, cluster coordinates) for significant differences were provided (0.5) | |
| 11. Conclusions were consistent with the results obtained and the limitations were discussed (1) | |
| | **TOTAL** 9.5 /11 |

Supplementary Table S2: Imaging Methodology Quality Assessment Checklist for 7.

"White matter integrity moderates the relation between experienced childhood maltreatment and fathers' behavioral response to infant crying" - Kim Alyousefi-van Dijk, Noa van der Knaap, Renate S M Buisman, Lisa I Horstman, Anna M Lotz, Madelon M E Riem, Carlo Schuengel, Marinus H van IJzendoorn, Marian J Bakermans-Kranenburg.

| **Category 1: Subjects** | **Score** (0/0.5/1) |
|---|---|
| 1. Patients were recruited prospectively and demographic data was reported (1) | |
| 2. Important variables (e.g. age, gender, race) were compared between the groups, either by stratification or statistically (1) | |
| 3. Sample size per group > 10 (1) | |
| **Category 2: Methods for image acquisition and analysis** | |
| 4. Magnet strength at least 1.5T (1) | |
| 5. MRI slice-thickness≤3 mm (1) | |
| 6. Whole brain analysis was automated with no a-priori regional selection (1) | |
| 7. Coordinates reported in a standard space (1) | |
| 8. The imaging technique used was clearly described so that it could be reproduced (1) | |
| 9. Measurements were clearly described so that they could be reproduced (1) | |
| **Category 3: Results and conclusions** | |
| 10. Statistical parameters (p-value, cluster size, cluster coordinates) for significant differences were provided (1) | |
| 11. Conclusions were consistent with the results obtained and the limitations were discussed (1) | |
| | **TOTAL** 11 /11 |

Bubb, E. J., Metzler-Baddeley, C., & Aggleton, J. P. (2018). The cingulum bundle: Anatomy, function, and dysfunction. *Neurosci Biobehav Rev*, *92*, 104-127. https://doi.org/10.1016/j.neubiorev.2018.05.008

Catani, M., Jones, D. K., Donato, R., & Ffytche, D. H. (2003). Occipito-temporal connections in the human brain. *Brain*, *126*(Pt 9), 2093-2107. https://doi.org/10.1093/brain/awg203

Costa, M., Braga, V. L., Yağmurlu, K., Centeno, R. S., Cavalheiro, S., & Chaddad-Neto, F. (2018). A Technical Guide for Fiber Tract Dissection of the Internal Capsule. *Turk Neurosurg*, *28*(6), 934-939. https://doi.org/10.5137/1019-5149.Jtn.20884-17.1

Emos, M., Khan Suheb, M., & Agarwal, S. ( 2024). *Neuroanatomy, Internal Capsule*. StatPearls Publishing. https://www.ncbi.nlm.nih.gov/books/NBK542181/

Frodl, T., Carballedo, A., Fagan, A. J., Lisiecka, D., Ferguson, Y., & Meaney, J. F. (2012). Effects of early-life adversity on white matter diffusivity changes in patients at risk for major depression. *J Psychiatry Neurosci*, *37*(1), 37-45. https://doi.org/10.1503/jpn.110028

George, K., & Das, J. (2024). *Neuroanatomy, Thalamocortical Radiations*. StatPearls Publishing. https://www.ncbi.nlm.nih.gov/books/NBK546699/

Goldstein, A., Covington, B., Mahabadi, N., & Mesfin, B. (2024). *Neuroanatomy, Corpus Callosum*. StatPearls Publishing. https://www.ncbi.nlm.nih.gov/books/NBK448209/

Janelle, F., Iorio-Morin, C., D'amour, S., & Fortin, D. (2022). Superior Longitudinal Fasciculus: A Review of the Anatomical Descriptions With Functional Correlates [Review]. *Frontiers in Neurology*, *13*. https://doi.org/10.3389/fneur.2022.794618

Jang, S. H. (2014). The corticospinal tract from the viewpoint of brain rehabilitation. *J Rehabil Med*, *46*(3), 193-199. https://doi.org/10.2340/16501977-1782

Javed, K., Reddy, V., & Forshing, L. (2024). *Neuroanatomy, Lateral Corticospinal Tract*. StatPearls Publishing. https://www.ncbi.nlm.nih.gov/books/NBK534818/

Patel, A., Biso, G., & Fowler, J. (2024). *Neuroanatomy, Temporal Lobe*. StatPearls Publishing. https://www.ncbi.nlm.nih.gov/books/NBK519512/

Rong, D., Zhang, M., Ma, Q., Lu, J., & Li, K. (2014). Corticospinal tract change during motor recovery in patients with medulla infarct: a diffusion tensor imaging study. *Biomed Res Int*, *2014*, 524096. https://doi.org/10.1155/2014/524096

Schmahmann, J. D., Pandya, D. N., Wang, R., Dai, G., D'Arceuil, H. E., de Crespigny, A. J., & Wedeen, V. J. (2007). Association fibre pathways of the brain: parallel observations from diffusion spectrum imaging and autoradiography. *Brain*, *130*(Pt 3), 630-653. https://doi.org/10.1093/brain/awl359

Von Der Heide, R. J., Skipper, L. M., Klobusicky, E., & Olson, I. R. (2013). Dissecting the uncinate fasciculus: disorders, controversies and a hypothesis. *Brain*, *136*(Pt 6), 1692-1707. https://doi.org/10.1093/brain/awt094

Wu, H., Melicher, T., Bauer, I. E., Sanches, M., & Soares, J. C. (2020). Chapter 3 - Brain Structural Abnormalities of Major Depressive Disorder. In R. S. McIntyre (Ed.), *Major Depressive Disorder* (pp. 39-49). Elsevier. https://doi.org/https://doi.org/10.1016/B978-0-323-58131-8.00003-3
